# Glycyrrhizin, an inhibitor of HMGB1 induces autolysosomal degradation function and inhibits *H. pylori* infection

**DOI:** 10.1101/2022.08.02.502584

**Authors:** Uzma Khan, Bipul Chandra Karmakar, Priyanka Basak, Sangita Paul, Animesh Gope, Deotima Sarkar, Asish Kumar Mukhopadhyay, Shanta Dutta, Sushmita Bhattacharya

**Affiliations:** ICMR-National Institute of Cholera and Enteric Diseases (ICMR-NICED), Kolkata-700010, India

## Abstract

*Helicobacter pylori* a key agent for causing gastric complications is linked with peptic ulcer, gastritis, and in severe cases gastric cancer. In response to infection, host cells stimulate autophagy to maintain cellular homeostasis. However, *H. pylori* have evolved the ability to usurp the host’s autophagic machinery. High mobility group box1 (HMGB1), an alarmin molecule is a regulator of autophagy and its expression is augmented in gastric cancer and many other cancers. Therefore, this study aims to explore the role of glycyrrhizin (a known inhibitor of HMGB1) in autophagy during *H. pylori* infection. Human gastric cancer (AGS) cells were infected with *H. pylori* SS1 strain and further treatment was done with glycyrrhizin. Western blot was used to examine the expression levels of autophagy proteins. Autophagy and lysosomal activity were monitored by immunofluorescence. We have performed knockdown of HMGB1 to verify the effect of glycyrrhizin by siRNA transfection method. *H. pylori*-infection *in vivo* C56BL/6 mice model was established and the effect of glycyrrhizin treatment was studied. We found that the autophagy-lysosomal pathway was impaired due to a significant increase in lysosomal membrane permeabilization during *H. pylori* infection in AGS cells. Subsequently, glycyrrhizin treatment restored the lysosomal membrane integrity, accompanied by an increase in cathepsin B activity and reduction of ROS and inflammatory cytokine IL-8. The recovered lysosomal function enhanced autolysosome formation and concomitantly attenuated the intracellular *H. pylori* growth by eliminating the pathogenic niche from gastric cells. Additionally, glycyrrhizin treatment inhibited inflammation and improved gastric tissue damages in mice.

## Introduction

Infection with *Helicobacter pylori* is one of the key factors responsible for causing gastric disorders and a major risk factor for progression to gastric ulcers and in severe forms gastric cancer (1–4). It is a gram-negative bacterium that has evolved with the ability to colonize and take refuge in epithelial and immune cells of the stomach (1–4). This is considered as one of the possible reasons owing to the rise of antibiotic resistance of *H. pylori* (5, 6). On account of this, WHO has laid down this organism into high-priority pathogens list (7). Mounting evidence suggests that reprogramming host cellular pathways are obligatory facets of *H. pylori* infection (8–11). On the other end of the spectrum, to eliminate an incoming pathogen, the host often deploys several cellular defence strategies. Among them, autophagy is one of the important pathways involved in recognizing and capturing intracellular bacteria for its degradation (10, 12–14). *H. pylori* infection in epithelial cells and macrophages often induces the host autophagic machinery during early infection while survival and colonization of *H. pylori* are favoured by inhibition of autophagy at later stages (14–17). Recent reports suggest that compromised autophagy benefits intracellular survival of *H pylori* (15). But, still, we are far from a definitive description of the molecular mechanisms through which circumvention of autophagy leads to the overall consequences in *H. pylori* pathogenesis.

Autophagy is a dynamic process that involves the double membrane-bound autophagosome formation which further fuses into the lysosome for degradation of autophagic substrates by lysosomal hydrolases (18, 19). Any harm incurred during the process results in impairment of autophagy (15, 17). Disrupted lysosomal function is one of the major causes of inactivation of autophagy during *H. pylori* infection. Studies have shown that damaged lysosomal function enables *H. pylori* intracellular survival within autophagosomes while invading gastric epithelial cells (13). To maintain its survival, *H. pylori* exploit the autophagy machinery with a precise mechanism. VacA protein secreted from *H. pylori* induces autophagosome formation at initial stages but impairs autophagic flux at later stages (13, 14). CagA another protein secreted is known to negatively regulate autophagy. This antagonistic behaviour of *H. pylori* secretory proteins facilitates its successful propagation within the autophagosome (19).

There are several host factors and proteins involved during *H. pylori* infection to regulate the autophagy mechanism. Autophagy protein SQSTM1/p62 degradation is inhibited during *H. pylori* infection for different time exposures (13, 18, 19). Further LAMP1 expression, one of the end-stage markers of autophagy is reduced due to *H. pylori* infection (13).Prior studies have shown that High mobility group box 1 (HMGB1) is augmented during *H. pylori* infection (20). Moreover, HMGB1 is a major player in different types of endocrine cancer as it modulates autophagy, apoptosis, and necrosis (21–24). HMGB1 interacts with RAGE to elicit cytokine response during infection (20, 25). It is known to interact with autophagy proteins like beclin1 and induce autophagy (24). Besides the fact that HMGB1 induces autophagy, recent reports are contradicting the phenomenon of HMGB1-induced autophagy (26). A growing line of evidence suggests that HMGB1 ruptures lysosomal membrane and initiates a chain of events including cathepsin B release into the cytosol (26, 27). Ruptured lysosomal membrane is the characteristics of lysosomal membrane permeabilization (LMP). This lysosomal dysfunction is also associated with *H. pylori* infection. Therefore, modifying HMGB1 production or preventing its interaction with other molecules might provide an opportunity for the prevention or treatment of gastric disorders.

The role of HMGB1 mediated autophagy in *H pylori* infection is unknown. Research over the past has established that HMGB1 modifies the host cell signalling system in gastric epithelial cells (29). It translocates in the cytosol and extracellular space during inflammation, trauma, necrosis, and oxidative stress (21-23, 28). Further, HMGB1 is considered an important link between autophagy and inflammation. Targeting the molecular mechanism of HMGB1 in autophagy during *H. pylori* infection might help us to develop effective treatment strategies for gastric disorders.

Keeping in mind this scenario of *H. pylori* mediated infection and gastric disorders; drug designing is inevitable as antibiotic resistance is well known. Treatment for *H. pylori* is complicated as successful eradication depends on several host and pathogenic factors. Triple therapy or sequential therapy is still one of the best ways of treatment but antimicrobial resistance has decreased the efficacy of these therapies (29, 30). An increase in failure of these therapies leads towards development of specific inhibitors. Recent reports showed that autophagy inducers like Vitamin D and statin decreased *H. pylori* burden (31, 32). Hence, identifying autophagy inducers can be an alternative therapeutic option for reducing *H. pylori* infection. As HMGB1 has a distinctive role in *H pylori* infection and autophagy, pharmacological inhibition or knockdown of HMGB1 might play an important part in reducing pathogenesis. In this study, we have used an inhibitor of HMGB1, glycyrrhizin to explore the role of the autophagy-lysosomal pathway during *H. pylori* infection in both *in vitro* and *in vivo* conditions.

## Materials and Methods

### *H. pylori* culture

*H. pylori*, Sydney Strain SS1 (cagA+, vacA s2m2) were grown on brain heart infusion (BHI) agar (Difco, USA) containing 7% heat-inactivated horse serum (Invitrogen), antibiotics and IsoVitaleX as mentioned previously (33). Plates were kept in a microaerophilic atmosphere at 37°C for five to six days. Stock cultures were stored at − 70°C for further usage. Isolates were re streaked on fresh BHI agar and incubated for 24 hours which was used for experimental studies. *H. pylori* resistant strain OT-14(3), (clarithromycin, metronidazole resistant) was cultured with the same protocol as of SS1.

### Cell Culture

Human gastric cancer cell line AGS was gifted by Dr. Asish Kumar Mukhopadhyay (ICMR-NICED, Kolkata). AGS cells, were grown in F12 media (Sigma-Aldrich) supplemented with 10% heat inactivated FBS (Sigma, USA), 1% penicillin-streptomycin (Sigma, USA) and maintained in an incubator at 37°C and 5% CO2.

### *In vitro* Infection Assay

Cell density of 0.5×10^6^ cell per 60mm cell culture dish was plated. *H. pylori* SS1 culture was dissolved in sterile phosphate buffered saline (PBS) and adjusted to an OD of 1 at 600 nm followed by centrifugation at 10,000 g for 10 min. The cells were starved overnight in antibiotic and FBS-free incomplete F12 media. Cells were further infected with or without *H. pylori* with a multiplicity of infection (MOI) 1:100 for 4 h followed by gentamicin treatment for 1 h, to kill the extracellular bacteria. Cells were then washed with PBS and incubated in fresh medium and treatment was done with glycyrrhizin GLZ (200μM) for another 4 h. Cell lysis was performed by adding 0.1% saponin for 15 min at room temperature and serial dilution was done before plating on BHIA plates to determine the invaded bacteria into the AGS cells. Colonies were then counted after 5-7 days culture (CFU/ml). Similar assay was also performed for *H. pylori* resistant strain [OT-14(3)] with or without glycyrrhizin for 4h. The cells and media supernatant were collected after infection for further experiments

### Real-Time PCR

RNA was isolated from infected and drug treated /siRNA transfected cells using RNA TRIzol reagent. Further, cDNA was prepared from RNA utilising Thermo Scientific c-DNAs synthesis kit. SYBR green kit of Applied Biosystems was used for Quantitative PCR. ΔΔCt method was used to calculate and normalization was performed with the housekeeping gene control GAPDH. ΔΔCt= test - internal control-test control. Relative density of *H. pylori* was quantified by performing semi-quantitative PCR, detecting *H. pylori*-specific 16S-ribosomal DNA (rDNA) primer, FP (5’-AGAGAAGCAATACTGTGAA-3′) & *H. pylori*, RP (5′-CGATTACTAGCGATTCCA-3′). GAPDH in the same samples were measured for normalization, FP 5′-GTCTTCACCACCATGGAGAAGGC-3′), and RP (5′-CATGCCAGTGAGCTTCCCGTTCA-3′).

### Immunofluorescence

AGS cells were seeded on coverslips and infection was done with *H. pylori* (4h) followed by glycyrrhizin incubation for 4h. For immunofluorescence staining, cells were fixed in 4% paraformaldehyde at room temperature for 1 h and blocked in PBS containing 3% BSA and 0.01% Triton X100 for 1 hr. Next, coverslips were incubated with anti-LAMP1 and anti-Galectin-3 at 4oC overnight. Subsequently, secondary antibody incubation was done using TRITC-conjugated anti-rabbit secondary antibody (1:1000) (Cat# AP132R) and FITC-conjugated anti-mouse secondary antibody (1:1000). Lastly, the coverslips were mounted on glass slides by adding ProLong™ Gold Antifade reagent with DAPI (Thermo Fisher) and examined using an inverted confocal microscope (Carl Zeiss LSM 710). For LAMP1 and LC3B double immunofluorescence staining same protocol was followed.

### Plasmid transfection

The tfLC3B plasmid was a gift from Dr. Dhiraj Kumar ICGEB, New Delhi, India. To examine autophagosomes and autolysosomes, AGS cells were transiently transfected with tf-LC3B plasmid using Lipofectamine 2000 (Invitrogen). 48 h after transfection, cells were incubated with or without *H. pylori* and treated with glycyrrhizin. At the end, coverslips were mounted on ProLong™ Gold Antifade reagent with DAPI (Thermo Fisher) and imaged using an inverted confocal microscope.

### Transient transfection

The following siRNAs were purchased from IDT: ATG5siRNA (ID hs.Ri. ATG5.13.1) and HMGB1 siRNA (ID hs.Ri.HMGB1.13.1) and used for transfection at 70 % confluence with siRNA/ Non-specific siRNA using lipofectamine in a 35mm dish. 48 h after transfection, cells were infected with or without *H. pylori* as described.

### Live-cell confocal microscopy

To monitor the lysosomes expressing LAMP1, *H. pylori* infected and drug treated AGS cells were subjected to live-cell imaging by adding 5μl of baculovirus expressing Lamp1-GFP construct (CellLight™ Lysosomes-GFP, BacMam 2.0,#C10507) and incubated for 16h then observed in confocal microscope.

### LysoTracker staining

AGS cells (2 × 105) were seeded on coverslips. *H. pylori* infection for 4 h was done after 24h followed by drug treatment. To investigate the acidification of lysosomes, cells were incubated with LysoTracker Red DND-99 (Invitrogen, L7528) for 30 min. Cell were then observed under an inverted confocal microscope.

### Magic Red staining

To monitor the activity of the lysosomal protease cathepsin B, Magic Red™ Cathepsin B substrate (red) staining was performed in *H. pylori* infected and drug treated AGS cells and imaged immediately under confocal microscope.

### Acridine Orange staining

Infection with *H. pylori* was performed for 4h in AGS cells grown in coverslips. After infection, cells were treated with glycyrrhizin for 4h. To evaluate lysosome membrane integrity, cells were stained with Acridine Orange (AO) staining for 30mins and observed under fluorescent microscope, Axio Observer7 Apotome.2.

### MTT Assay

Cellular toxicity was examined using a Colorimetric Cell Viability Kit (MTT) (Promokine) in 96-well plates. MTT reagent was added and kept for 4 h. Purple crystal formazan formed was solubilised with DMSO. Amount of formazan salt was measured in a microplate reader (Bio-Rad Serial no. 19901) at OD of 590 nm.

### ROS levels

Intracellular ROS levels were monitored by using 2,7-dichlorodihydrofluorescein diacetate (DCFH-DA). 10μM of DCFH-DA was added to control, infected and drug treated cells for 30 min and kept at 37°C. Excess DCFH-DA was washed with PBS three times. Finally, fluorescence was monitored (Ex-485 nm and Em-520 nm) using a multimode reader, Molecular devices Spectramax M2.

### Immunoblotting

Control, treated and infected cells were lysed in RIPA (Radio immuno precipitation assay buffer) lysis buffer containing protease and phosphatase inhibitors. After cell lysis centrifugation was done at 7000 rpm for 20 mins at 4°C. Further, protein level was determined and run on 10% or 12.5% SDS-PAGE gel at 120V. Gels were further transferred to PVDF membrane. 5% skimmed milk dissolved in TBST (20mMTris-HCl, 150 mM NaCl, 0.1% Tween20) buffer was used for blocking and incubated for 1h at room temperature. In the next step, the membranes were kept overnight with primary antibodies at 4°C. The primary antibodies used are: rabbit polyclonal Anti-SQSTM1/P62 antibody (Cat# ab91526), rabbit monoclonal anti-ATG5 (Cat #ab228668), mouse monoclonal anti-LAMP1 (Cat# 15665S), rabbit monoclonal anti-beclin1 antibody (Cat #ab207612), rabbit polyclonal anti-LC3B antibody (Cat#ab51520), mouse polyclonal anti-β-actin (Cat #sc-47778), mouse monoclonal anti-Galectin-3 (Cat #sc-53127), rabbit polyclonal anti-α-Tubulin (Cat #BB-AB0118), anti-rabbit secondary HRP-conjugate (Cat #12-348), anti-mouse secondary HRP-conjugate (Cat #12-349).

### Enzyme-linked immunosorbent assay (ELISA)

Pro-inflammatory cytokine (IL-8, IL-6) levels from media and serum were estimated using Krishgen Biosystems kit as per manufacturer’s instructions. All experiments were done in triplicate.

### *H. pylori* infection in C57BL/6 mice and treatment with Glycyrrhizin

Mice were maintained in the animal house under 12-hour dark light cycles. Experiments were conducted under the guidelines of Institutional Animal Ethical Committee, NICED, Kolkata (PRO/157/-260 July 2022). 8 weeks of C57BL/6 mice bred in house were used for the experiments. Three different experimental set of mice were grouped: Control group (CON), n = 4, *H. pylori* SS1 infected group (HP), n = 4, infected group treated with GLZ (HP+GLZ), n = 4. All groups of mice were treated every day for seven days with antibiotic cocktail (Ciprofloxacin, Metronidazole, Erythromycin, Albendazole) to avoid any other bacterial/parasites infections. Group of mice (HP & HP+GLZ) were inoculated with 10^8^ CFU/mouse/inoculation of SS1 on three alternative days, or PBS (CON). After two weeks of inoculation, group of mice (HP+GLZ) were orally injected with glycyrrhizin (10 mg/kg) for 4 weeks, while group of mice (CON) received sterile water. At the end of week 4, all mice were sacrificed. Gastric tissues were isolated and blood was collected for experimental purpose.

### Statistical Analysis

All data were represented as mean ± S.E.M. Two groups were compared using Unpaired t-test, and multiple comparisons were done by one-way ANOVA. Significance level has been marked as, * for p < 0.05, which implies significant, ** for p < 0.01, which implies very significant, *** for p < 0.001, which implies highly significant.

## Results

### Glycyrrhizin induces autophagy in gastric epithelial cells

Previous reports indicated that glycyrrhizin induces autophagy in myoblast cells but there are no such reports in gastric cells till date (34). Here we checked the expression levels of different autophagy proteins upon glycyrrhizin treatment in AGS gastric cancer cells. Glycyrrhizin treatment for 4h elevated the expression of autophagy proteins LC3IIB, LAMP1, and beclin1 in a dose-dependent manner. Similarly glycyrrhizin inhibited HMGB1 expression at different concentrations (**Fig. 1A**). The most effective dose was 200μM. (**Fig. 1A**). To examine the possibility of toxicity of glycyrrhizin, we checked the effect of glycyrrhizin on the viability of AGS cells (**Fig. S1A**). Glycyrrhizin treatment for 24h at different concentrations (50, 100, 200μM) did not show significant toxicity. Further, we confirmed glycyrrhizin-induced autophagy by immunofluorescence of LC3IIB. Drug treatment for 4h showed an enhancement of LC3IIB puncta formation significantly (**Fig. 1B**). Subsequently, we assessed the effect of glycyrrhizin induced autophagosomal maturation in gastric cancer cells. LAMP1 is known to be a marker for lysosomal activity, therefore, we labelled lysosomes with LAMP1-GFP construct after exposure to glycyrrhizin treatment. Subsequently, glycyrrhizin induced LAMP1 expression (green) significantly in live cells than the control (**Fig. 1C**). Together these results suggest that glycyrrhizin induces an autophagic response in gastric cells.

**FIG 1.**
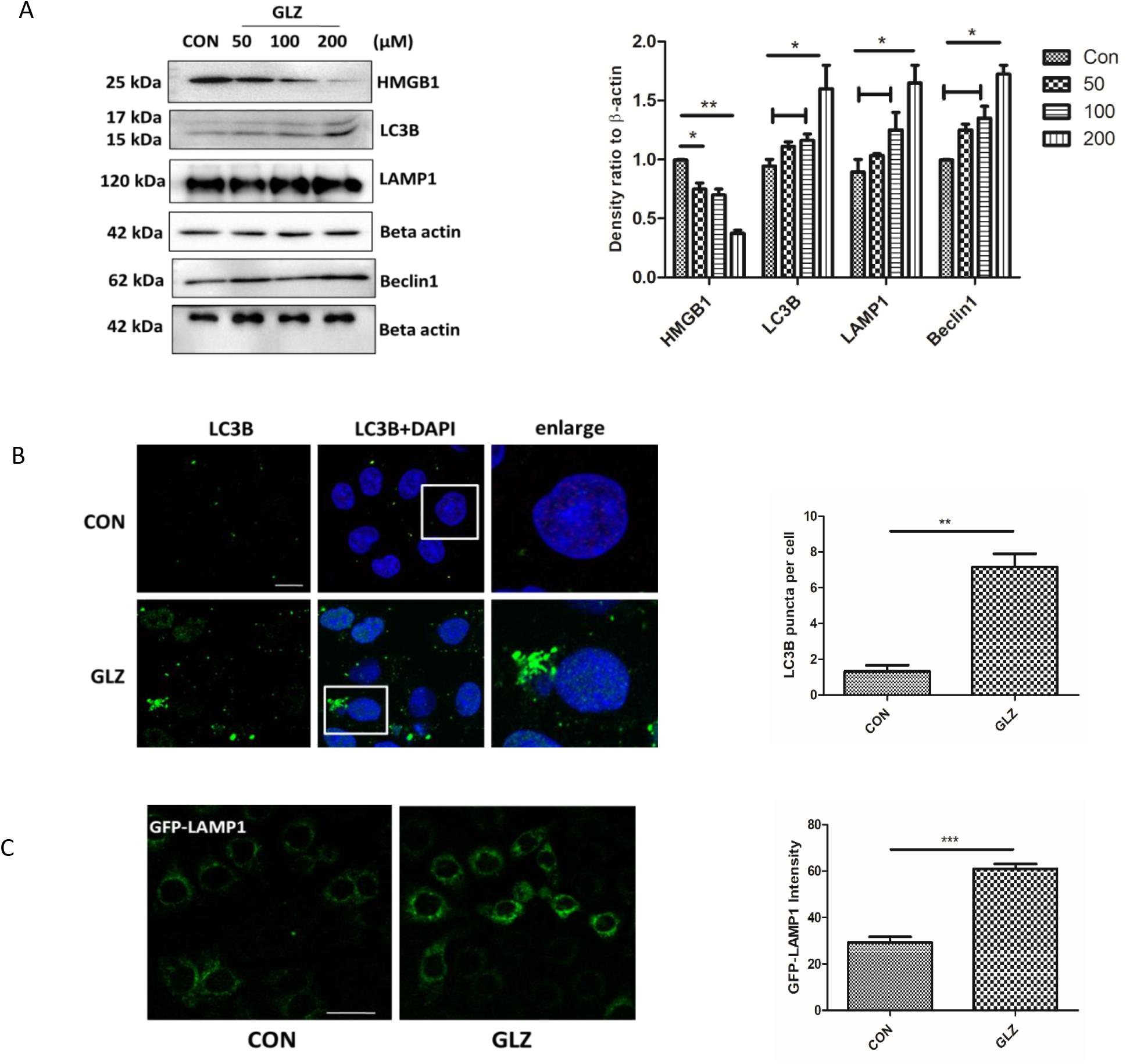
Glycyrrhizin treatment overexpresses autophagy proteins. AGS cells were treated with glycyrrhizin GLZ (200μM) or DMSO for 4 h (A) Cell lysates were prepared and expression level of HMGB1 and autophagy marker proteins LC3B, LAMP1, and Beclin1 was observed by western blot analysis. Beta-actin was used as protein loading control. Densitometric analyses are represented graphically. (B) Control and drug treated cells were subjected to immunofluorescence. Scale bar: 5μm. Confocal microscopy showed LC3B puncta (green) formation. LC3B puncta formation was quantified and graphically plotted. (C) Live cell imaging using LAMP1-GFP construct for labelling lysosomes was performed and we examined under confocal microscopy, GFP-LAMP1 fluorescence intensity was measured. Scale bar: 10μm. Graphs were represented as mean±SEM (n=3); Unpaired t-test was done and significance was calculated; * p < 0.05, ** p < 0.01, *** p < 0.001

### Glycyrrhizin induced autophagy inhibits intracellular *H. pylori* growth

Since *H. pylori* is known to invade gastric epithelial cells, we examined the expression of autophagy proteins by immunoblotting in *H. pylori* infected gastric cancer cells. *H. pylori* Sydney Strain SS1 was used for infection in gastric cells for 4 h and post treatment was done with glycyrrhizin (4h). We observed upregulation of LC3IIB and LAMP1 in glycyrrhizin-treated *H. pylori*-infected cells (**Fig. 2A**). Moreover, LC3IIB and LAMP1 were also upregulated in glycyrrhizin-treated control cells as previously explained in (**Fig. 1A, B**). Glycyrrhizin driven autophagy induction also enhanced p62 expression in infected and uninfected cells as compared to only infected cells. To verify the findings of LC3IIB and LAMP1 expression, we additionally performed an immunofluorescence assay and live-cell analysis of drug treated and *H. pylori* infected cells. Consistently, an enhanced LC3B puncta formation and an increased LAMP1 expression were observed upon glycyrrhizin exposure as compared to untreated infected cells (**Fig. 2B, C**). The results showed that autophagosomal and lysosomal activities are increased by glycyrrhizin.

**FIG 2.**
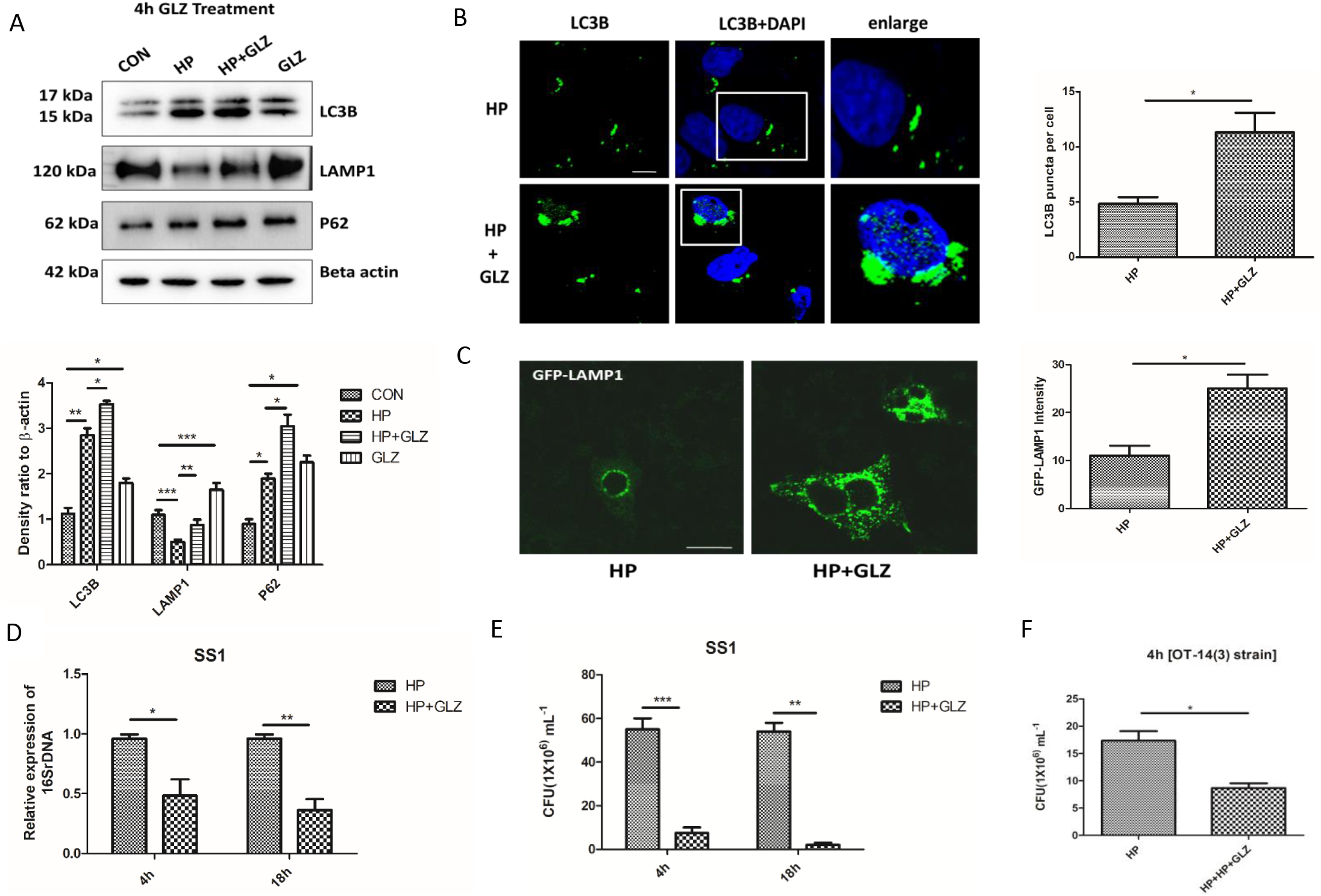
Exposure with Glycyrrhizin reduces intracellular *H. pylori* growth in AGS cells. (A-C) Infection with *H. pylori* SS1 strain (MOI 100) was performed in cells for 4 h and further exposed to glycyrrhizin (GLZ) (200μM) for 4 h. (A) Immunoblotting was performed for quantification of autophagy-associated marker proteins (LC3B, LAMP1, and P62). Beta-actin was used as loading control. Densitometry analyses are represented graphically. One-way ANOVA was performed. (B) Confocal microscopy showed LC3B puncta (green) in *H. pylori* (HP) infected & *H. pylori+* glycyrrhizin (HP+GLZ) treated cells after immunofluorescence LC3B puncta formation was quantified and graphically plotted. Scale bar: 5μm. (C) Live cell imaging of LAMP1 under confocal microscopy showed GFP-LAMP1 puncta formation. Unpaired t-test was performed and graphically represented Scale bar: 10μm. (D-E) Cells were incubated with *H. pylori* SS1 strain (MOI 100) for 4 h followed by gentamicin treatment to kill extracellular bacteria. Finally, cells were treated with glycyrrhizin GLZ (200μM) at two different time points for 4h and 18h (D) Intracellular *H. pylori* DNA (16SrDNA) was determined by real-time PCR. GAPDH was used as the internal control. (E) Cells were lysed and plated on BHIA plates with serial dilutions, for 4-5 days for counting colony and CFU/ml was graphically represented. (F) Infection with *H. pylori* resistant strain (OT-14(3) (MOI 100) for 4 h was performed in gastric cells followed by GLZ (200μM) treatment for 4h and CFU/ml was graphically represented. Graph were represented as mean±SEM (n=3); Unpaired t-test was done and significance was calculated; * p < 0.05, ** p < 0.01, *** p < 0.001

Next, we sought to examine the effect of autophagy induction on intracellular bacterial growth. We performed real-time PCR (RT-PCR) and checked *H. pylori-specific* 16SrDNA. Intracellular *H. pylori* level was significantly reduced by glycyrrhizin treatment for 4h and 18h (**Fig. 2D**). Of interest, we additionally examine the bacterial proliferation by bacterial adhesion assay. In line, intracellular *H. pylori* burden was decreased significantly due to drug exposure for 4h and 18h (**Fig. 2E**). Since antimicrobial resistance is a problem to curb *H. pylori* infection, we treated a resistant strain of *H. pylori* with glycyrrhizin for 4h in AGS cells. Glycyrrhizin significantly reduced the growth of resistant *H. pylori* (OT-14 (3)) strain (**Fig. 2F**). However, glycyrrhizin failed to reduce *H. pylori* growth in BHIA media which indicates glycyrrhizin has no direct bactericidal effect on *H. pylori* at 200μM concentration (**Fig. S2A**). Taken together, these data indicate that glycyrrhizin induces autophagy in gastric cancer cells and inhibits intracellular *H. pylori* growth.

### Enhancement of autophagic flux by glycyrrhizin contributed to anti-*H. pylori* activity

As *H. pylori* infection is involved in defective autosomal lysosomal maturation and degradation, we determined the effect of glycyrrhizin in lysosomal degradation activities. Hence, we checked the level of SQSTM1/p62, a key protein involved in autophagy. Western blotting in the previous figure revealed that at 4h, p62 is increased due to *H. pylori* infection whereas glycyrrhizin reversed the effect. Further, we observed that long term exposure of *H. pylori*, resulted in p62 accumulation at 18h. This is evident due to the impairment of autolysosomal degradation. On the other hand, glycyrrhizin treatment induces p62 degradation at 18h significantly, which is a hallmark of induction of lysosomal degradation (**Fig. 3A**). Further to assess the activation of autophagic flux by glycyrrhizin we performed a double immunofluorescence assay for both LC3B and LAMP1 protein. Results demonstrated that both LC3B and LAMP1 colocalized in *H. pylori*-infected and glycyrrhizin-treated infected and uninfected cells (**Fig. 3B**). The data indicated that glycyrrhizin induced autophagosomal lysosomal maturation. Next, we examined the stage of glycyrrhizin-mediated autophagic degradation in both autophagosomes and lysosomes by transfecting the AGS cells with tandem fluorescent LC3B (tfLC3B) plasmid. Autophagosomes appeared as yellow dots due to colocalization of both GFP and RFP. This has been observed in *H. pylori*-infected gastric cells whereas in glycyrrhizin-treated infected cells we observed more free red dots as GFP and RFP did not colocalize. Autolysosomes appears red due to the acidic pH of lysosomes which quench GFP (**Fig. 3C**). Taken together these results revealed that glycyrrhizin eliminates the *H. pylori*-mediated prevention of autolysosomal formation in the infected cells by promoting lysosomal maturation and subsequent destruction.

**FIG 3.**
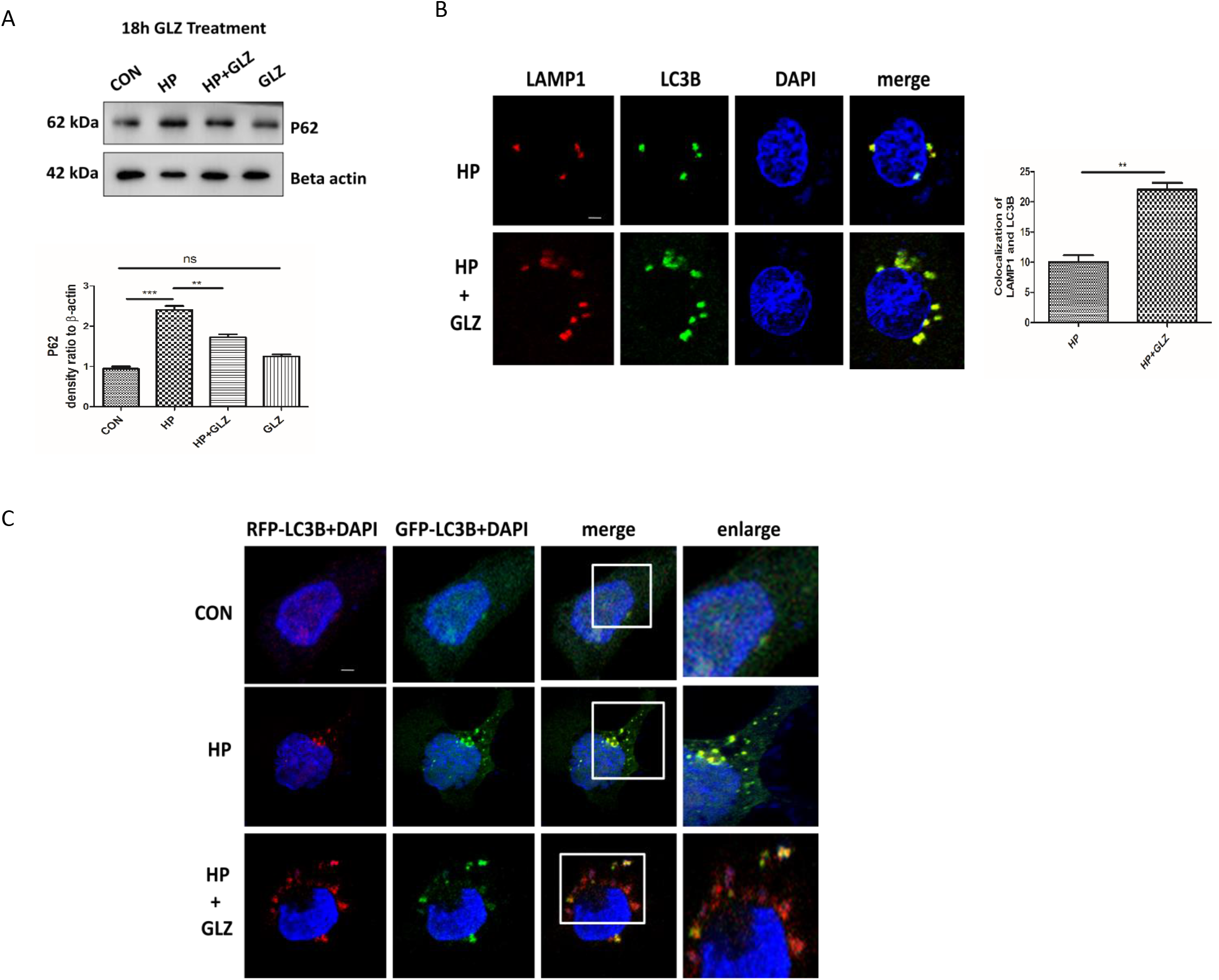
Activation of autophagy by glycyrrhizin contributes to anti-*H. pylori* activity. (A) Cells were incubated with *H. pylori* SS1 strain followed by glycyrrhizin GLZ (200μM) for 18h. Immunoblot analysis was performed for p62 protein. Beta-actin was used as loading control. Densitometry analyses are represented graphically. One-way ANOVA was performed. (B) Glycyrrhizin (GLZ) exposure to *H. pylori*-infected cells for 4 h was subjected to LAMP1 and LC3B double immunofluorescence. Confocal microscopy showed yellow puncta as colocalization of LC3B puncta (green) & LAMP1 (red). Colocalization intensity is graphically represented. Scale bar: 2μm. (C) AGS cells were transfected with a tandem mRFP-GFP tag (tfLC3B) plasmid & further infected with *H. pylori* SS1 *strain* (MOI 100) for 4 h and finally glycyrrhizin (GLZ) (200μM) treatment was done for 4h. Confocal microscopy showed LC3B puncta formation in control (CON), H *pylori* (HP)infected & *H pylori* + glycyrrhizin (HP+GLZ) treated cells. The yellow puncta showed autophagosomes. The free red puncta are autolysosomes. Scale bar: 2μm. Graph represented as mean±SEM (n=3); Significance was determined by Unpaired t-test; * p < 0.05, ** p < 0.01, *** p < 0.001

### Anti-*H. pylori* effect and autophagic degradation by glycyrrhizin is mediated through HMGB1 inhibition

To confirm or rule out the possible involvement of HMGB1 in *H. pylori* infection, AGS cells were transiently transfected with non-specific siRNA and HMGB1-specific siRNA followed by infection with *H. pylori* for 4h and subjected to immunoblotting. Western blot revealed that HMGB1 is silenced in AGS cells after 48h transfection (**Fig. S3A**). HMGB1 knockdown elevated the level of LAMP1 and LC3IIB significantly (**Fig. 4A**). In addition, we examined the level of intracellular *H. pylori* by RT-PCR. HMGB1 silencing attenuated intracellular *H. pylori* burden significantly (**Fig. 4B**).

**FIG 4.**
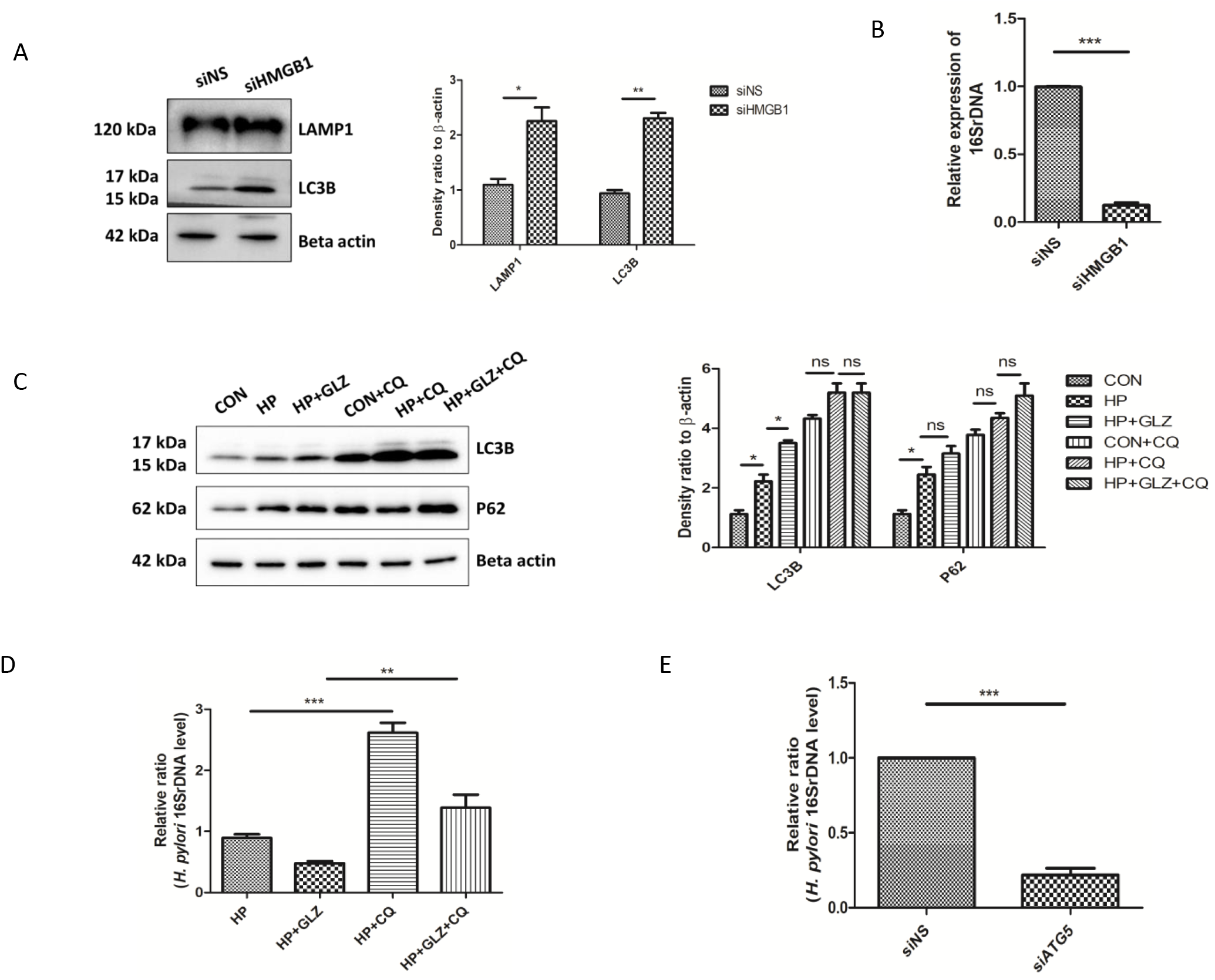
HMGB1 inhibition reduces bacterial growth while impairment of lysosomal activity induces bacterial survivability. (A-B) After transfection for 48h, scrambled siRNA (siNS) and HMGB1 siRNA (siHMGB1) transfected cells were further infected with *H. pylori* SS1 strain (MOI of 100) for 4 h. (A) Immunoblotting was performed in infected cell lysates for expression of LAMP1 and LC3B. Beta-actin was used as protein loading control. Densitometry analyses are represented graphically. (B) Intracellular *H. pylori* DNA (16SrDNA) were determined by RT-PCR. GAPDH was used as the internal control. Unpaired t-test was performed (c-d) AGS cells incubated with the *H. pylori* SS1 strain were exposed to glycyrrhizin GLZ (200μM) for 4h and/or chloroquine CQ (50 μM) for 4h. (C) Cell lysates were subjected to western blot to determine LC3B and P62 protein levels. Beta-actin was used as protein loading control. Densitometry analyses are graphically represented (D) Intracellular *H. pylori* DNA (16SrDNA) was measured by real-time PCR. GAPDH was used as the internal control. (E) Cells were transfected with non-specific siRNA (siNS) and ATG5 siRNA (siATG5) & then incubated with *H. pylori* SS1 strain for 4 h, intracellular *H. pylori* DNA was measured by RT-PCR. GAPDH was kept as an internal control. Graphs were represented as mean±SEM (n=3); One-way ANOVA was performed and significance was calculated; * p < 0.05, ** p < 0.01, *** p < 0.001

To validate the effect of glycyrrhizin on autolysosomal activity, we treated the AGS cells with chloroquine (CQ), a potent lysosomal inhibitor. LC3IIB accumulation and P62 aggregates increased in both *H. pylori*-infected and uninfected cells treated with glycyrrhizin after exposure to CQ (**Fig. 4C**). Additionally, we analyzed the effect of CQ in glycyrrhizin-mediated *H. pylori* clearance. Due to blocking of autophagic flux, CQ promptly elevated *H. pylori* infection and counteracted the antimicrobial action of glycyrrhizin (**Fig. 4D**). Our results demonstrated that chloroquine treatment inhibited anti-*H. pylori* activity of glycyrrhizin due to impaired autolysosomal degradation function.

According to previous reports, *H. pylori* can survive within non-digestive autophagosomes (13). We analyzed the effect of Atg5 knockdown (Atg5 is an autophagy marker protein required for autophagosome formation) on intracellular *H. pylori* survival. Western blot showed that Atg5 is silenced in AGS cells after 48h transfection (**Fig. S3B**). Consistently, bacterial clearance occurred significantly due to the silencing of Atg5 (**Fig. 4E**).

Cumulatively, these data demonstrated that *H. pylori* usurp autophagosomes for its perpetuation inside the host while glycyrrhizin, an inhibitor of HMGB1 promote autophagosomal lysosomal degradation which in turn reduces intracellular *H. pylori* growth.

### Lysosomal acidification was recovered by glycyrrhizin

Further, we investigated in detail the restored lysosomal degradation capacity of glycyrrhizin. We applied Acridine Orange staining to monitor the acidic compartment of lysosomes in AGS cells. Lower orange intensity in *H. pylori*-infected cells was observed as compared to control because lysosomal acidification was compromised whereas glycyrrhizin treatment increased orange intensity by restoring lysosomal acidification (**Fig. 5A**). Additionally, to evaluate lysosomal acidification, we exposed the cells to Lysotracker Red which selectively binds to vesicles that have low pH. Here, *H. pylori* infection affected lysosomal pH and reduced the fluorescent signals as compared to control. While, glycyrrhizin exposure restored the acidic pH and showed red signals in infected cells as compared to only infected cells (**Fig. 5B**). The data indicates that glycyrrhizin restored the disrupted lysosomal function during *H. pylori*-infection.

**FIG 5.**
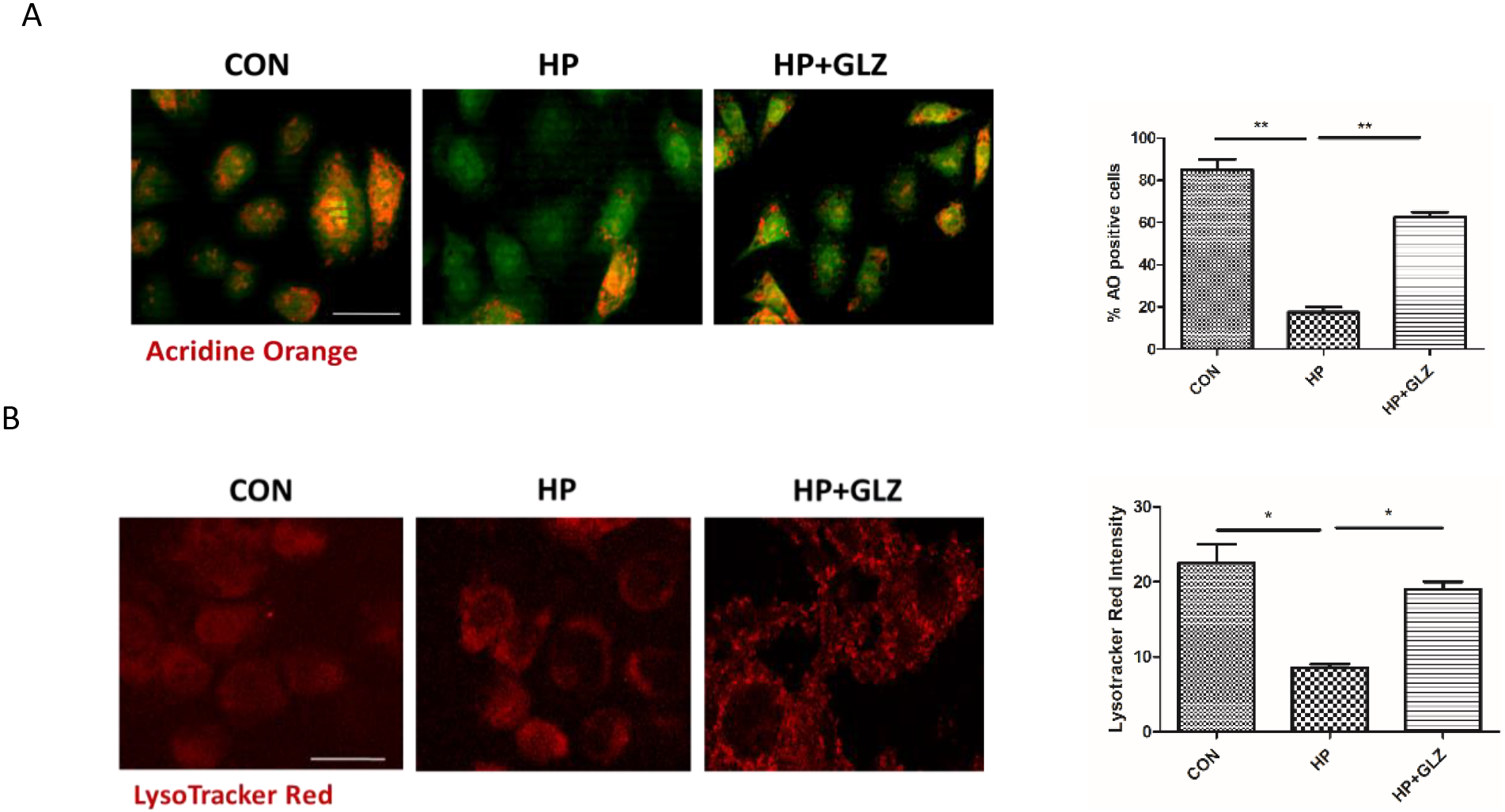
Lysosomal function is restored by glycyrrhizin. (A-B) Infection with *H. pylori* SS1 strain (MOI 100) was performed for 4 h followed by glycyrrhizin (GLZ) (200μM) exposure for 4h. (A) Lysosomal membrane integrity was monitored by Acridine Orange (AO) staining in a fluorescence microscope. Briefly, cells were incubated with 10μg/ml of acridine orange (15 min) and examined. Percentage of positively stained cells were measured and graphically represented. Scale bar: 10μm. (B) Drug treated, infected and control cells were incubated with LysoTracker Red (100 nM, 30 min) to label lysosomes and examined under the confocal microscope. Red fluorescent intensity was quantified. Scale bar: 10μm. Graphs were generated using GraphPad Prism 5 and represented as mean±SEM (n=3); Significance was calculated by one-way ANOVA; * p < 0.05, ** p < 0.01, *** p < 0.001

### Glycyrrhizin enhanced lysosomal degradation by inhibiting LMP

According to previous reports, *H. pylori* triggered undigested autophagosome accumulation resulting in lysosomal membrane permeabilization (LMP) (45). Moreover, HMGB1 is also involved in LMP (26, 27). Therefore, we determined whether the effect of glycyrrhizin in restoring lysosomal function is mediated by LMP. We performed double immuno-fluorescence of galectin3 and LAMP1 (LAMP is exposed to glycoproteins of lysosomal membrane after LMP). Colocalization of galectin3 with LAMP1 remarkably increased in *H. pylori* infection than control as galectin3 binds to lysosomal membrane glycoproteins which are exposed after LMP. But, glycyrrhizin treatment in infected cells reduced co-localization of galectin3 and LAMP1 probably due to inhibition of LMP (**Fig. 6A**). Lysosomal proteases such as cathepsins are activated and released into the cytosol during LMP. Hence, we further determined cathepsin B activity by Magic Red staining. Glycyrrhizin treatment significantly increased red fluorescent staining as compared to *H. pylori*-infected cells (**Fig. 6B**). This reveals that glycyrrhizin is able to reduce LMP by restoration of the lysosomal acidification and integrity of the lysosomal membrane. Subsequently, we examined the effect of inhibition of LMP by glycyrrhizin on ROS and inflammatory cytokines as LMP is linked to inflammation and oxidative stress. In line, glycyrrhizin exposed cells dramatically reduced ROS level and IL-8 secretion significantly as compared to only infected cells (**Fig. 6C, D**). Taken together these data indicate that glycyrrhizin improved lysosomal degradation activity and induced protective effects by inhibiting LMP.

**FIG 6.**
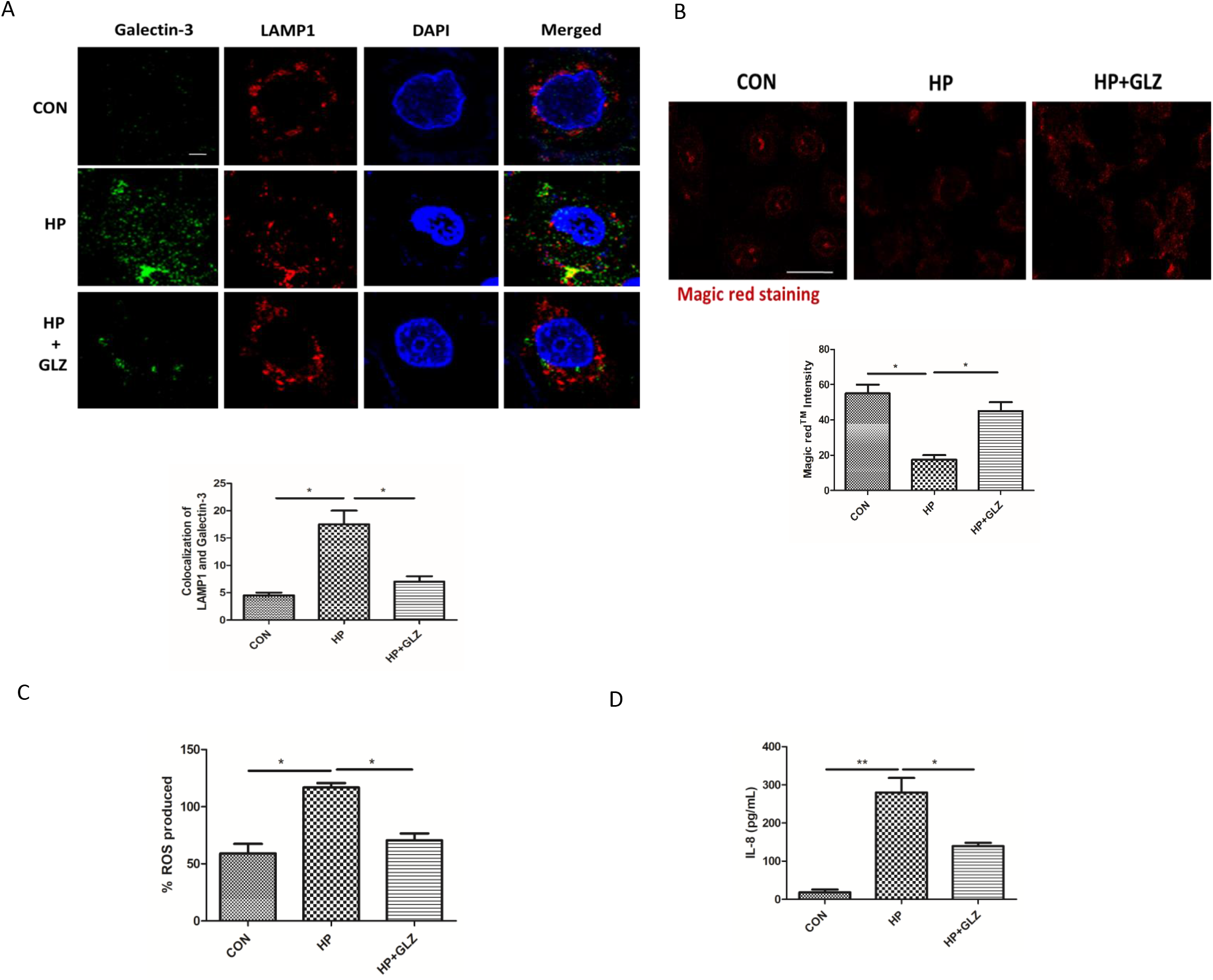
Lysosomal membrane integrity is restored by glycyrrhizin. (a-d) Infection with *H. pylori* SS1 strain (MOI 100) was performed for 4 h followed by glycyrrhizin (GLZ) (200μM) exposure for 4h. (A) Double immunofluorescence was done with LAMP1 & Galectin-3 antibodies to confirm LMP. Cells were observed under the confocal microscope and colocalization intensity was measured. Scale bar: 2μm. (B) Cathepsin B activity was examined after staining with Magic Red™ for 1 h and imaging was done under the confocal microscope. Scale bar: 10μm. Fluorescence intensity was measured. (C) Expression of reactive oxygen species (ROS) in infected and drug treated cells was determined by DCFDA methods for 30min in a fluorimeter. (D) The expression of IL-8 from media collected after treatment was analysed by ELISA in a microplate reader. Graphs were generated using GraphPad Prism 5 and represented as mean±SEM (n=3); Significance was calculated by one-way ANOVA; * p < 0.05, ** p < 0.01, *** p

### In *in vivo* mice model, glycyrrhizin induces autophagy and reduces gastric damages

Eventually, we validated the activity of glycyrrhizin in *in vivo* mice model. We have infected mice with *H. pylori* SSI strain. After infection, mice were treated with glycyrrhizin for 30 days at a 10mg/kg body weight dose (**Fig. S4A**). Effective dose of glycyrrhizin was determined for *H pylori* infected mice from previous data of glycyrrhizin (34, 35). We performed western blots with control, infected and glycyrrhizin treated infected gastric tissues. Glycyrrhizin reduced the level of HMGB1 significantly and induced p62 degradation which is a marker of autophagosomal lysosomal degradation. It also augmented LAMP1 expression (**Fig. 7A**). However, Glycyrrhizin reduced inflammation and inhibited epithelial cell damages. Moreover, we checked the effect of glycyrrhizin on IL-6 levels in the serum collected from treated mice. Glycyrrhizin significantly reduced IL-6 expression (**Fig. 7B**). To further assess the anti-*H. pylori* effect of glycyrrhizin, we examined gastric tissues for changes in the morphology. There were changes in the gastric histopathology when compared with a control group for *H. pylori*-infected gastric tissues (**Fig.7C**). *H. pylori* infection-induced inflammation and inflammatory cell infiltration in gastric tissues have caused epithelial cell damage. Hence, collectively these data revealed that glycyrrhizin induced autophagy and consequently decreased inflammation and gastric tissue damages.

**FIG 7.**
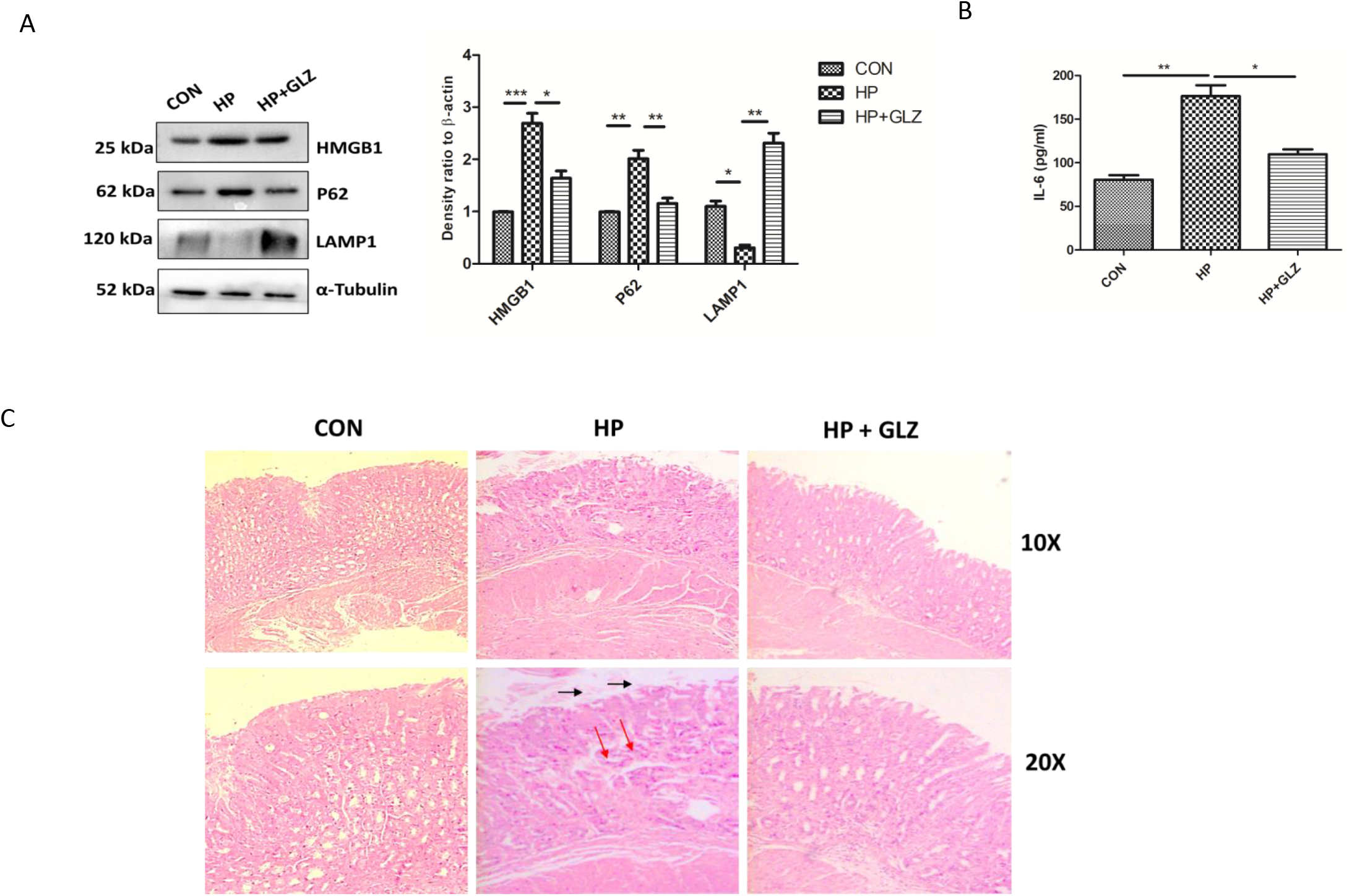
Glycyrrhizin treatment induces autophagy in *H. pylori*-infected mice and ameliorates gastric tissue damages (A-D) C57BL/6 mice (n = 4 per group) were treated with antibiotics every 7 days. Then after 7 days incubation period, mice were infected with *H. pylori* SS1 strain thrice a week on alternate days. Mice were incubated for 14 days and then administered with or without glycyrrhizin GLZ (10 mg /kg body weight), every other day for 4 weeks. After treatment, mice were sacrificed, gastric tissues and serum were collected. (A) Immunoblot showing the expression of HMGB1 and autophagy proteins (HMGB1, P62, LAMP1) of mouse gastric tissues. α-tubulin was used as protein loading control. Densitometry analyses are represented graphically. (B) The expression of IL-6 was determined by ELISA in a microplate reader. (C) Histology images of Control (CON*), H. pylori* (HP) infected, and *H. pylori*-infected plus glycyrrhizin treated (HP+GLZ) gastric tissues at 10X and 20X respectively representing the inflammatory changes. Black arrows (↑) indicate stomach gastric epithelium, red arrows (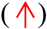) indicate gastric damage. Graphs were represented as mean±SEM (n=3); Significance was determined by one-way ANOVA; * p < 0.05, ** p < 0.01, *** p < 0.001

## Discussion

Current evidence suggests that autophagy plays a major role to protect the host from bacterial pathogens (36, 37). But the pathogens have their own mechanisms to subvert autophagy and persistently invade the host and promote intracellular survival. *Salmonella, Shigella*, and *Mycobacterium* are known to avoid autophagy (38–41). However, there are reports which showed that *H. pylori* induces autophagy at the beginning but gradually it inhibits autophagy (13, 15). There is always a crosstalk between lysosomes and autophagosomes during bacterial pathogenesis to maintain a steady autophagic flux. The mechanisms behind initial activation and subsequent impairment involve a complex interplay between host and bacterial factors. *H. pylori* secrete virulent factors like CagA and VacA (13). Both these factors contribute towards pathogenesis. Although previous literature suggests that Vac A induces autophagy but later it has been reported by Raju et al that prolonged exposure to Vac A inhibits autophagy ((13, 42)). Recent studies have also revealed that autophagy is down-regulated in CagA+ strains as compared to CagA mutant strains (43). This inhibition of autophagy is accompanied by accumulation of p62 and decreased LAMP1 expression (43). Henceforth, in the current study, we have used *H. pylori* SS1 strain which is CagA positive but expresses non-functional VacA and this strain is also capable of mice infection.

Here, we investigated the effect of an autophagy inducer, glycyrrhizin in inhibiting *H. pylori* infection. Glycyrrhizin is an inhibitor of HMGB1 which plays a major role in infectious diseases and cancer (20, 34, 44). HMGB1 is also reported to be overexpressed in *H. pylori*-infected gastric cells. Recent reports suggested that HMGB1 induces autophagy but at the same time also causes impairment of autophagy by inducing lysosomal membrane permeabilization (LMP) in diabetic retinopathy (26). Hence, we have targeted HMGB1 for treating *H. pylori* infection. Here, glycyrrhizin, an inhibitor of HMGB1 decreased the intracellular *H. pylori* burden in gastric cancer cells. To find out the details behind bacterial clearance, we observed that glycyrrhizin induces autophagy in gastric cells. This is consistent with previous studies in myoblast cells (34). Glycyrrhizin treatment attenuated *H pylori* infection and also induced expression of autophagy marker proteins. Moreover, glycyrrhizin showed co-localization of both LC3B and LAMP1 in *H. pylori*-infected gastric cancer cells. We also observed that LAMP1 expression has increased due to glycyrrhizin treatment. This indicates autolysosome formation as LAMP1 expression is necessary for autophagosomal maturation. Moreover, keeping in mind that growing antibiotic resistance of *H. pylori* is a major problem, glycyrrhizin was tested for its ability to eliminate the resistant *H. pylori* strain *in vitro*. Glycyrrhizin successfully showed clearance of antibiotic resistant *H. pylori*. In addition, transient knock down of HMGB1 resulted in parallel to glycyrrhizin treatment, hence the probable mechanism behind the induction of autophagy by glycyrrhizin is due to its inherent anti-HMGB1 property. Bacterial clearance by autophagy induction is a general mechanism as recent reports revealed that autophagy inducers like vitamin D and statin controlled *H. pylori* infection (31, 32). Moreover, in the past, researchers have shown that *H. pylori* infection inhibits lysosomal degradation by inducing lysosomal membrane permeabilization (45). Hence, we determined the effect of glycyrrhizin in the later stages of autophagy. Consistent with the results of initial autophagy induction, our results further demonstrated that glycyrrhizin treatment augments lysosomal degradation resulting in reduced bacterial burden. As it is reported that *H. pylori* induces p62 accumulation at later stages we performed 18h of infection. In line, we proved that p62 accumulation takes place and on the other hand, glycyrrhizin augmented p62 degradation. Subsequently, LC3IIB and p62 accumulation was observed using a lysosomal inhibitor chloroquine CQ. Glycyrrhizin further induces LC3IIB accumulation due to CQ treatment and survivability of *H pylori* indicating the involvement of autophagic flux during infection (31).

To gain deeper insight into the reason behind the lysosomal degradation of glycyrrhizin we searched for possible mechanisms. Previous studies suggested that *H. pylori* survive in undigested autophagosomes (13). Consistently, we proved that Atg5 knockdown inhibited *H. pylori* growth as Atg5 is responsible for autophagosome formation. Furthermore, the accumulation of undigested autophagosomes results in lysosomal membrane permeabilisation (LMP) (45). Moreover, HMGB1 is also associated with LMP via the Cathepsin B-dependent pathway and *H pylori* is also reported to reduce cathepsin activity due to disruption of lysosomal acidification (26, 45). In this study, inhibition of HMGB1 by glycyrrhizin rescued LMP in *H. pylori* infected cells and further restored the degradative capacity of autophagy proving that LMP is responsible for inhibition of autophagosome degradation. Moreover, we observed that lysosomal pH has been restored by glycyrrhizin using Lysotracker Red staining as acidic pH is extremely important for the proper digestive action of lysosomes. Further, we examined Acridine Orange and Magic Red activity to understand the mechanism of inhibition of LMP by glycyrrhizin. Here, also we found that glycyrrhizin exposure induced cathepsin B activity. Cathepsins are released in cytosol from lysosomes and show impaired activity during LMP (46). Additionally, HMGB1 and *H pylori* infection are responsible for decreased cathepsin B activity. Thus it has been proved that glycyrrhizin induces autophagy and lysosomal degradation by reducing LMP. Moreover, LMP is linked to the activation of ROS and inflammation (47). We evaluated the effect on ROS generation and cytokine expression. ROS generation and inflammatory cytokine expression are commonly enhanced during infection (48, 49). Consistently, the results showed that glycyrrhizin inhibited ROS production and inflammatory cytokine level in AGS cells. Therefore, our findings confirmed that inhibiting HMGB1 by glycyrrhizin induced autophagosomal maturation and lysosomal degradation in *H. pylori*-infected gastric cells by rescuing from LMP.

Further, we verified our findings *in vivo* mice model. In line, our results showed inhibition of HMGB1 expression and induction of autophagy by glycyrrhizin in gastric tissues. Moreover, histology and ELISA studies revealed that reduction of inflammation and recovery of gastric tissue damages due to glycyrrhizin treatment. Inflammation is a major problem during *H. pylori* infection. Inflammation induces gastric damages and causes further complications Here, Glycyrrhizin induces autophagy accompanied with reduction in inflammation. Hence, glycyrrhizin can be a potential drug candidate in near future to treat *H. pylori* infection and gastric disorders.

## Conclusion

In summary, our results demonstrated for the first time that induction of autophagy by inhibiting HMGB1 can reduce *H. pylori* infection in both *in vitro* and *in vivo* conditions. In addition, our data revealed that restoring the degradative capacity of autophagy by inhibiting HMGB1-induced LMP resulted in inhibition of *H. pylori* pathogenesis (**Fig. 8**). As both the host and pathogen play critical roles in disease progression, induction of autophagy and lysosomal degradation further affects downstream responses like inflammation and ROS generation. Hence, in the future glycyrrhizin might be used as a potent inducer of autophagy that reduces *H. pylori*-infection to inhibit progression of gastric disorders. The mechanism of antibacterial action of glycyrrhizin would provide novel strategies and targets to address the problem of antimicrobial resistance. Glycyrrhizin could also be used in synergistic composition with other drugs for *H. pylori* infection as standard *H. pylori* treatment requires triple therapy.

**FIG 8.**
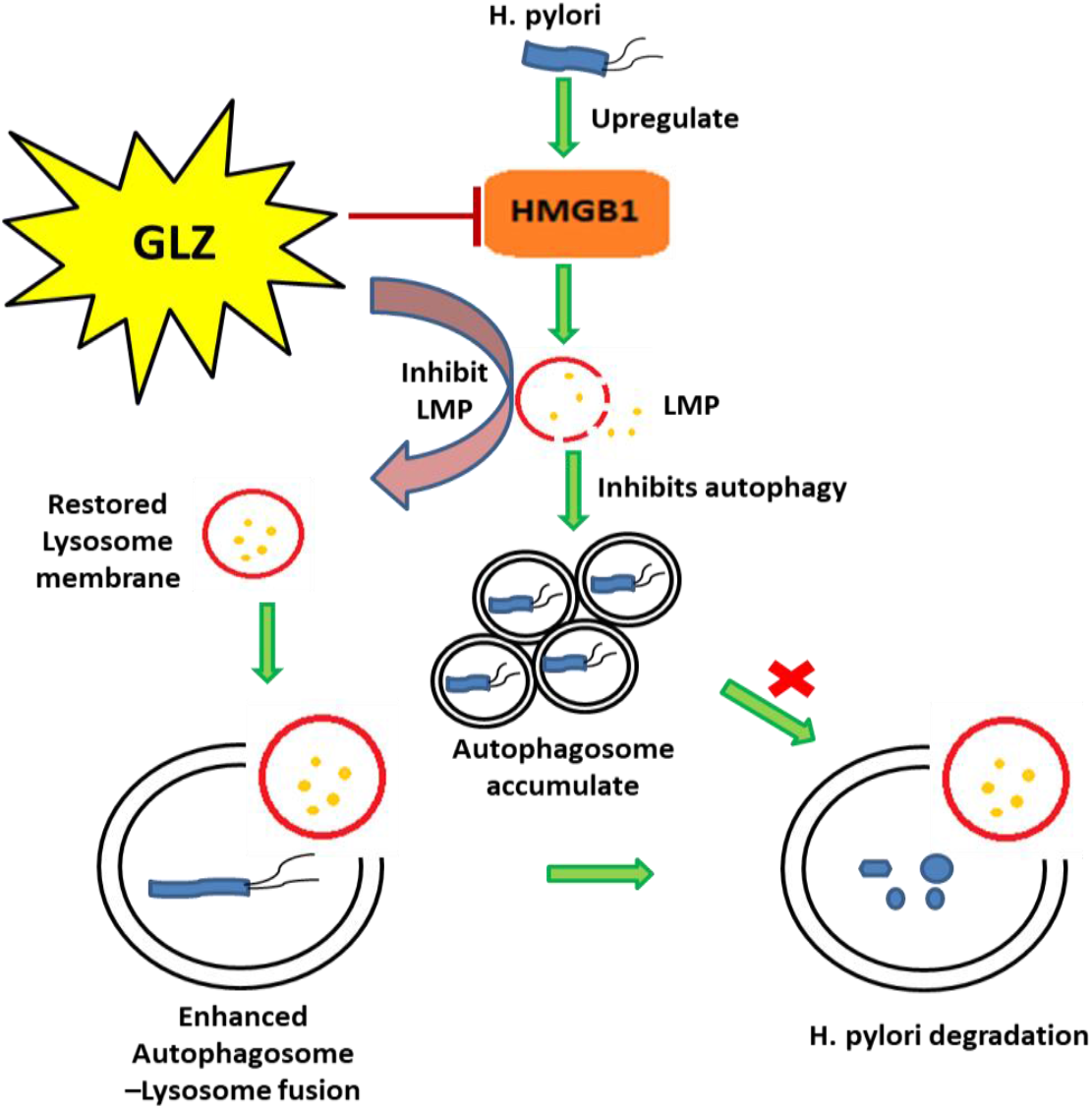
Schematic diagram representing the mechanism by which glycyrrhizin induces autophagy through inhibiting lysosomal membrane permeabilization (LMP). Glycyrrhizin inhibits HMGB1 which subsequently reduces LMP and induces autophagosomal lysosomal fusion to degrade *H. pylori*.

## Acknowledgements

This research was funded by a grant from Department of Biotechnology (DBT), Government of India, and Project (BT/PR3093/BIC/101/1076/2018). Authors thank Indian Council of Medical Research (ICMR), Council for Scientific and Industrial Research (CSIR), Department of Biotechnology (DBT) AND University Grants Commission (UGC) for fellowship assistance.

The authors declare no conflict of interest.

